# Biofoundry-scale DNA assembly validation using cost-effective high-throughput long read sequencing

**DOI:** 10.1101/2023.09.19.558498

**Authors:** Peter Vegh, Sophie Donovan, Susan Rosser, Giovanni Stracquadanio, Rennos Fragkoudis

**Affiliations:** Edinburgh Genome Foundry, School of Biological Sciences, University of Edinburgh

**Keywords:** biofoundry, DNA assembly, plasmid validation, sequencing

## Abstract

Biofoundries are automated high-throughput facilities specialising in the design, construction and testing of engineered/synthetic DNA constructs (plasmids), often from genetic parts. A critical step of this process is assessing the fidelity of the assembled DNA construct to the desired design. Current methods utilised for this purpose are restriction digest or PCR followed by fragment analysis, and sequencing. The Edinburgh Genome Foundry (EGF) has recently established a single-molecule sequencing quality control step using the Oxford Nanopore sequencing technology, along with a companion Nextflow pipeline and a Python package to perform in-depth analysis and generate a detailed report. Our software enables biofoundry scientists and end-users to rapidly analyse sequencing data, without specialised bioinformatics knowledge. In conclusion, we have created a laboratory and software protocol that validates assembled, cloned or edited plasmids, using Nanopore long reads, which can serve as a useful resource for the genetics, synthetic biology and sequencing communities.

**Author information:** All authors contributed to the design of the sequencing quality control step and pipeline, and the preparation of the manuscript. P.V. wrote the manuscript, designed and implemented the bioinformatics pipeline and interpreted results. S.D. wrote the manuscript, implemented the laboratory protocol and interpreted results. G.S. designed the bioinformatics pipeline. R.F. wrote the manuscript and contributed to the design of the laboratory protocol and pipeline.

Address: Edinburgh Genome Foundry (University of Edinburgh), Michael Swann Building, Max Born Crescent, Edinburgh, EH9 3BF, United Kingdom

**Graphical abstract:** 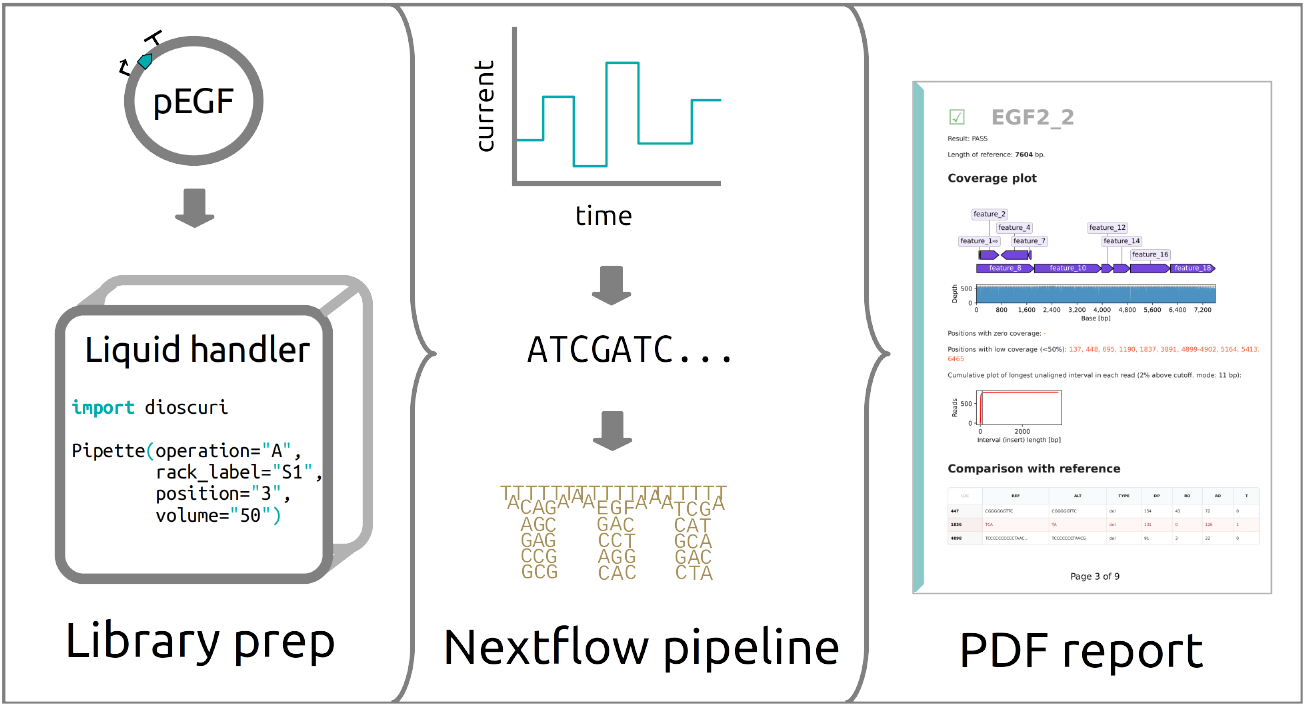

## Introduction

A critical step of the Design Build Test Learn (DBTL) cycle widely adopted in synthetic biology^1^ is verifying the fidelity of the DNA construct, obtained in the Build phase, to the designed DNA sequence. Several factors may lead to erroneous DNA constructs, including incorrect input DNA, problems during assembly or sample handling, misannealing overhangs, addition of point mutations, and homologous or other forms of recombination. Currently, the verification step relies mostly on restriction digest or PCR followed by fragment analysis (FA), and sequencing^2^. FA provides a cost-efficient but indirect, low confidence confirmation of construct correctness by checking the presence of specific restriction enzyme recognition sites and the fragment size.

Conversely, DNA sequencing approaches provide a nucleotide-level readout at the expense of a substantially higher cost. Sanger sequencing, for example, is not feasible in most cases due to cost and high number of reactions, but it may be useful for verifying targeted regions or in small batches of similar constructs assembled from shared genetic parts^2^. Second and thirdgeneration sequencing methods provide a solution to sequencing large batches of plasmid constructs due to their high-throughput and no requirement for using primers. In this technical note, we describe a single-molecule sequencing DNA assembly quality control solution at the Edinburgh Genome Foundry (EGF), that can be utilised by biologists and the sequencing community. EGF is an automated high-throughput facility (biofoundry) specialising in the modular assembly of DNA constructs (plasmids), using Golden Gate cloning. EGF’s platform is species agnostic and its outputs are used in projects as diverse as programming of stem cells for personalised medicine applications, vaccine development, gene therapy and many more.

## Results and Discussion

The DNA verification by sequencing step at the end of the Build phase of a synthetic biology (engineering) project aims at comparing the fidelity of an assembled DNA construct to its corresponding designed sequence. Here we present a one-step software pipeline to interpret sequencing data, which requires no specialised bioinformatics knowledge (Figure 1). Specifically, we wanted to obtain an annotated comparison of the sequenced and the expected DNA, and a judgement call for each sample, based on various checks.

**Figure 1.**
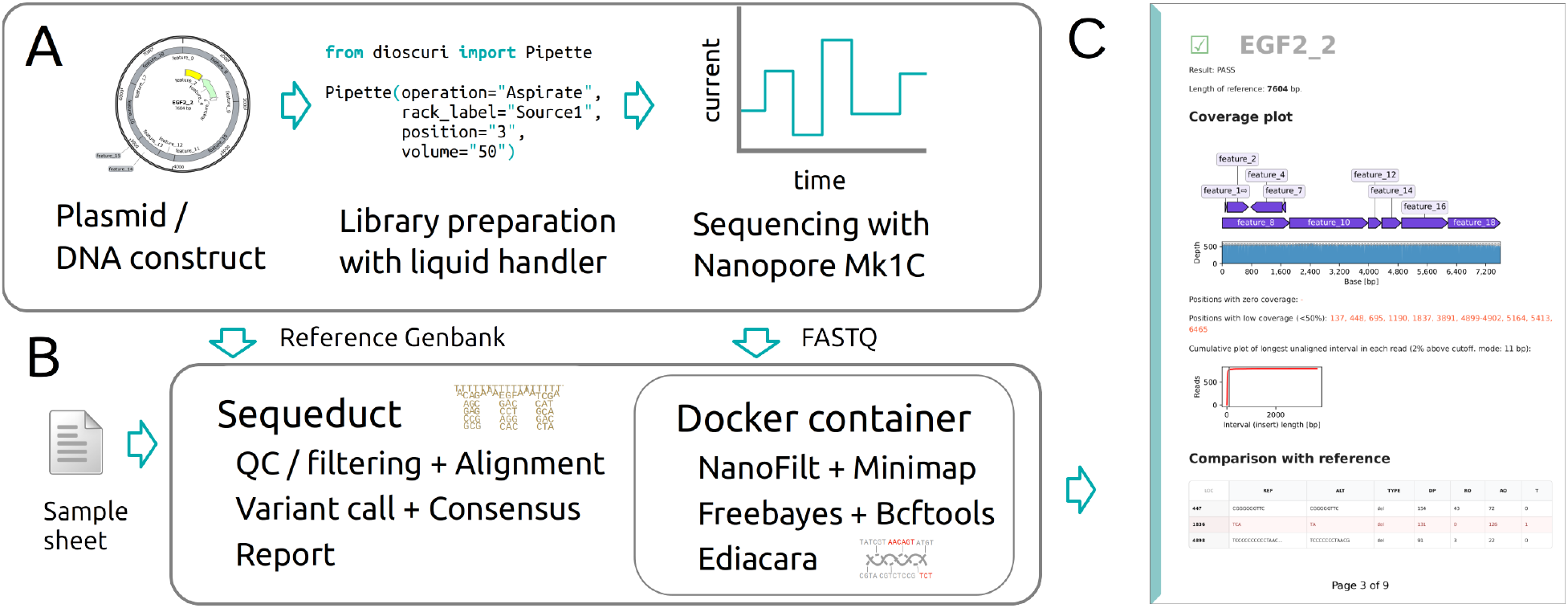
An overview of the sequencing pipeline. (A) The assembled or cloned plasmids are prepared into libraries, using a liquid handling platform. Libraries are loaded onto a Flongle flow cell in an Oxford Nanopore Mk1C sequencer. (B) The FASTQ files are analysed with a Nextflow pipeline (Sequeduct) that utilises a Docker container with all required software. (C) The Ediacara Python package creates a PDF report for an easy overview and interpretation of the results.

We hereby refer to *errors* as differences (or mutations, variants) between the designed and assembled DNA sequence. These errors include single nucleotide variants (SNV), small insertions and deletions (indels), structural variations (SV), and sequencing errors^3^. Although random sequencing errors can be mitigated with increased sequencing depth, systematic errors are much harder to avoid; here we focused on SNVs and SVs.

We apply automated protocols for the Oxford Nanopore Technologies (ONT) rapid barcoding kits to simultaneously fragment each sample (plasmid or DNA construct) and ligate individual barcodes. Up to 96 barcoded samples are pooled into a single library and loaded onto Flongle flow cells in a MinION Mk1C sequencer (see Methods section). A Nextflow^4^ pipeline, named Sequeduct, has been created to perform alignment, variant detection and reporting (https://github.com/Edinburgh-Genome-Foundry/Sequeduct/). This pipeline requires the FASTQ files of the partial and full length reads obtained from sequencing, the reference Genbank files of the designed sequences, and a sample sheet. The generated PDF report contains a chapter for each barcode: read statistics are followed by a histogram of the read lengths, which is a good indicator of the presence of structural errors. A displayed coverage chart with the reference sequence visualizes any deletions (Supporting information S1). Visualizing insertions in a similar plot is not straightforward, therefore a cumulative plot of the longest unaligned interval is provided as an indicator of large insertions. A variant (error) table is also provided in a simplified variant call format (VCF)^5^, which lists point and small variants. Variants at homopolymer stretches are flagged as this is a known systemic error^3^. Variants are also annotated on the reference sequence on a second plot, for an easy overview and navigation. Based on the above results, each of the plasmids with sufficient number of reads are assigned a pass/fail outcome, as detailed in Methods. The pipeline also returns the consensus FASTA sequence for each barcode.

If structural variations are found, then a subsequent task is to describe its nature and provide an explanation. This is largely beyond the scope of validation, but a second pipeline is also provided which aligns the genetic part sequences against the variant call consensus – or a *de novo* plasmid sequence assembled with Canu^6^ – and reports and visualizes the alignments in a PDF file (Supporting information S2). This is useful for evaluating plasmids that are constructed from parts using various toolkits, such as EMMA^7^, MoClo^8^ or Mobius^9^, to clarify whether we have part or sample mix-ups, recombination events or overhang misannealing.

Multiple alternative approaches have been published by Oxford Nanopore Technologies and other research laboratories. The EPI2ME Clone validation workflow uses *de novo* assembly to produce a FASTA file for each sample^10^. The SequenceGenie workflow analyses data from a multiplexed sequencing approach using a novel sample barcoding system^11^. The MinION Plasmid Sequence Verification Pipeline provides a cost-effective way of sequencing plasmids for clinical research applications^12^. Circuit-seq creates *de novo* assemblies from multiplexed samples^13^ while OnRamp is reference-based^14^. In comparison to these, Sequeduct performs an evaluation against an expected sequence and focuses on the produced report and downstream interpretation of results, more suitable for engineering biology and quality control purposes.

Several companies provide Nanopore sequencing service of plasmids for a fee (∼$15 / plasmid). Performing the sequencing in-house is cost-competitive and fast, provided that at least 24 samples are sequenced at the same time. In any case, Sequeduct is free software, and can be used with FASTQ data from sequencing providers in order to generate a more detailed and targeted report.

Ongoing development aims to incorporate additional functionalities and improvements. These include deconvoluting mixed samples, where multiple plasmids are in the same sample or use the same barcode. This would allow sequencing polyclonal (“polyploid”) samples or combinatorial assemblies, or pooling multiple plasmids into the same barcode. Similarly, we plan to address more error-scenarios in the pipeline as we accumulate sequencing results. We anticipate that analysis of synthetic design outcomes using full length sequence results will lead to the establishment of more robust DNA design rules.

In conclusion, we have set up a complete solution – consisting of laboratory and software protocols – for the validation of assembled, cloned or edited plasmids, using long reads. The software is available under a free and open-source license (GPLv3) to encourage contributions and feedback from biofoundries and the sequencing community.

## Methods

### Sequencing

Plasmid DNA is prepared using the Wizard SV 96 Plasmid DNA Purification System (Promega) and the concentration is measured by fluorescence-based quantification (Qubit dsDNA BR Assay Kit, Thermo Scientific). Samples are normalised to be within the 20 – 90 fmol/μL range and 1 μL of sample is used for library preparation. The protocols for the ONT Rapid Barcoding Kit (SQK-RBK004) or Rapid Barcoding Kit 96 (SQK-RBK110.96) are performed following the manufacturer’s instructions on an Opentrons OT-2 (SQK-RBK004) or Tecan Freedom EVO200 (SQK-RBK110.96) liquid handling robot. Libraries are loaded onto Flongle flow cells (R9.4.1) and run for up to 24-hours on a MinION Mk1C device that performs basecalling using Guppy v4.3.4.

### Analysis

The ‘pass’ folder of the FASTQ sequencing data is used in the analysis. The pipeline is written in Nextflow and is available on GitHub with documentation and an example dataset at https://github.com/Edinburgh-Genome-Foundry/Sequeduct. The first workflow (‘preview’) generates summary plots of each barcode for an overview of the sequencing run, using NanoPlot, part of the NanoPack suite^15^. The second workflow (‘analysis’) filters FASTQ files using NanoFilt followed by read alignment to the reference sequence using minimap2^16^. The reference sequence files can be created with a sequence editor or batch simulated using DNA Cauldron^17^. SAMtools^18^ is used to obtain coverage data and variants are called with freebayes^19^. Consensus sequence files are created with BCFtools^20^. As part of the pipeline, a Python package, Ediacara, was also written to generate a report PDF that visualises results for each barcode / plasmid (https://github.com/Edinburgh-Genome-Foundry/Ediacara). The package utilises Biopython^21^ and DNA Features Viewer^22^. Each plasmid is assigned an outcome: samples with insufficient number of reads (sequencing problems) resulting below 30x coverage are marked as ‘low coverage’. Remaining samples are marked ‘fail’ if problems are detected: zero coverage sections, majority of reads having an unaligned (insert) segment, consensus sequence length outside tolerance. The ‘warning’ label is applied for cases where the errors are below the set threshold levels. All other samples are marked as ‘pass’. An explanation of the report is also included in the Appendix section of the report. In addition to the pipeline, a Dockerfile is also provided to generate a Docker image with all required software.

## Acknowledgement

The Edinburgh Genome Foundry is supported by the BBSRC (BB/M018040/1) and the BBSRC/MRC/EPSRC-funded UK Centre for Mammalian Synthetic Biology as part of the RCUK’s Synthetic Biology for Growth program. This work was supported by the UKRI EPSRC Fellowship (EP/V033794/1) to G.S.

## Abbreviations

VCF: variant call format
DBTL: design build test learn
FA: fragment analysis

## Supporting information

Supporting information S1: example PDF report of the ‘analysis’ pipeline.

Supporting information S2: example PDF report of the ‘review’ pipeline.

