## Supporting information S1 for "Biofoundry-scale DNA assembly validation using cost-effective high-throughput long read sequencing"

### Sequencing analysis report

This document reports the sequencing results of the **EGF demo** project. There are **2** barcodes in the analysis. Each chapter details the results of one barcode, and within a chapter each section reports on one plasmid. Please see the Appendix on the last page for an explanation of the report.

There are **1623** filtered reads in the analysed barcodes of this sequencing run.

### Barcode: barcode01

Results for the **1** construct(s) in the pool: **1** / **0** / **0** / **0** (pass / warning / fail / low coverage).

| Name | Result | Length [bp] | FASTQ reads | Coverage [x] |
|---|---|---|---|---|
| **EGF2_2** | PASS | 7604 | 792 | 557 |

Histogram of the **792** FASTQ reads:

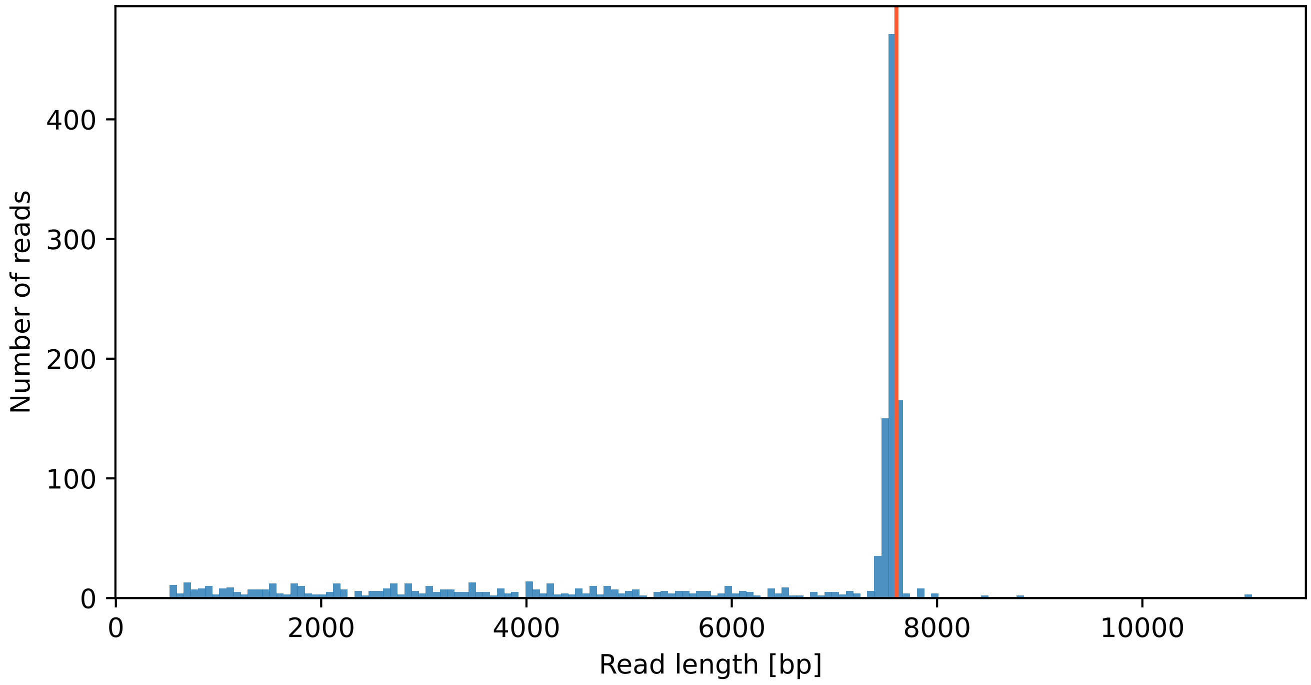


The vertical red lines show the expected construct lengths.

### ☑ EGF2_2

Result: PASS

Length of reference: **7604** bp.

#### Coverage plot

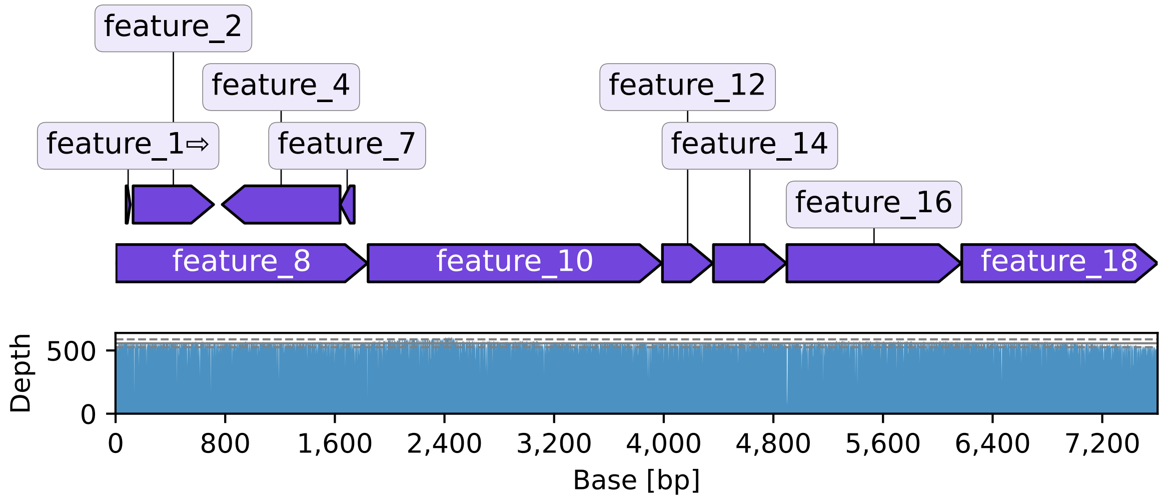


Positions with zero coverage: -

Positions with low coverage (<50%): 137, 448, 695, 1190, 1837, 3891, 4899-4902, 5164, 5413, 6465

Cumulative plot of longest unaligned interval in each read (2% above cutoff. mode: 11 bp):

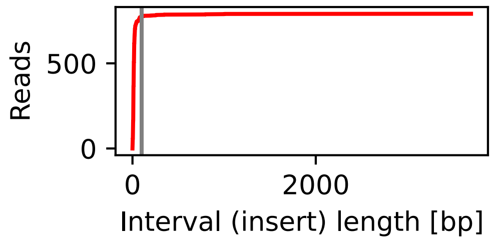


#### Comparison with reference

| LOC | REF | ALT | TYPE | DP | RO | AO | T |
|---|---|---|---|---|---|---|---|
| **447** | CGGGGGGTTC | CGGGGGTTC | del | 154 | 43 | 72 | 0 |
| **1836** | TCA | TA | del | 131 | 0 | 126 | 1 |
| **4898** | TCCCCCCCCCCTAAC... | TCCCCCCCTAACG | del | 91 | 3 | 22 | 0 |

**EGF2_2** reference vs consensus of reads:

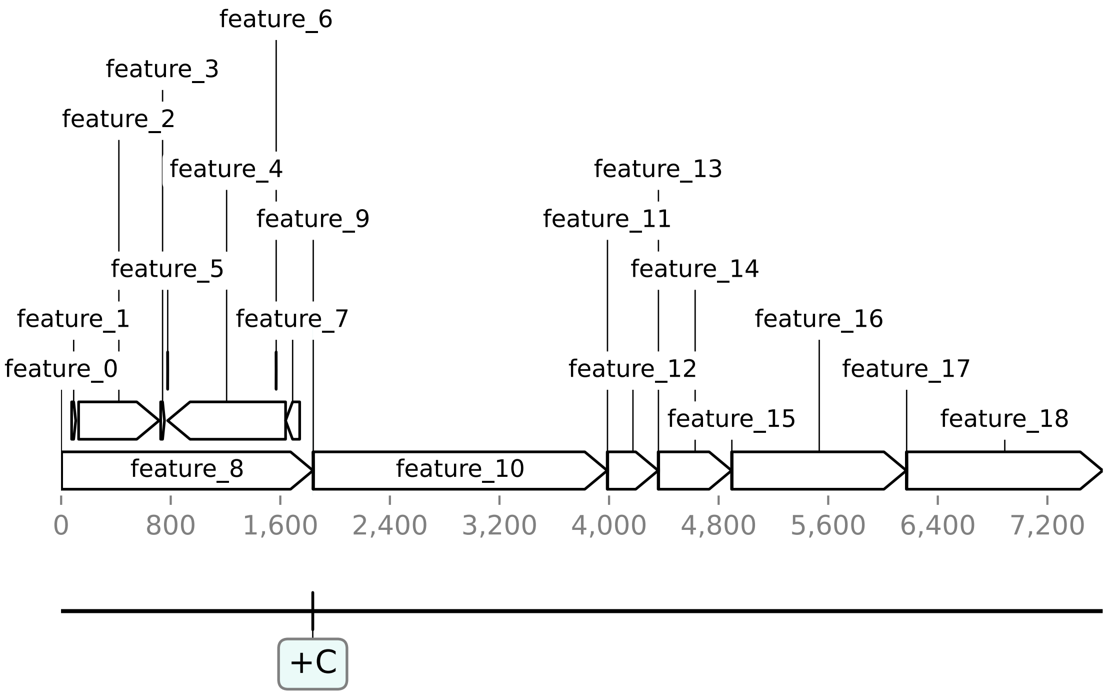

### Barcode: barcode12

Results for the **1** construct(s) in the pool: **0** / **0** / **1** / **0** (pass / warning / fail / low coverage).

| Name | Result | Length [bp] | FASTQ reads | Coverage [x] |
|---|---|---|---|---|
| **EGF2_13** | FAIL | 8939 | 831 | 592 |

Histogram of the **831** FASTQ reads:

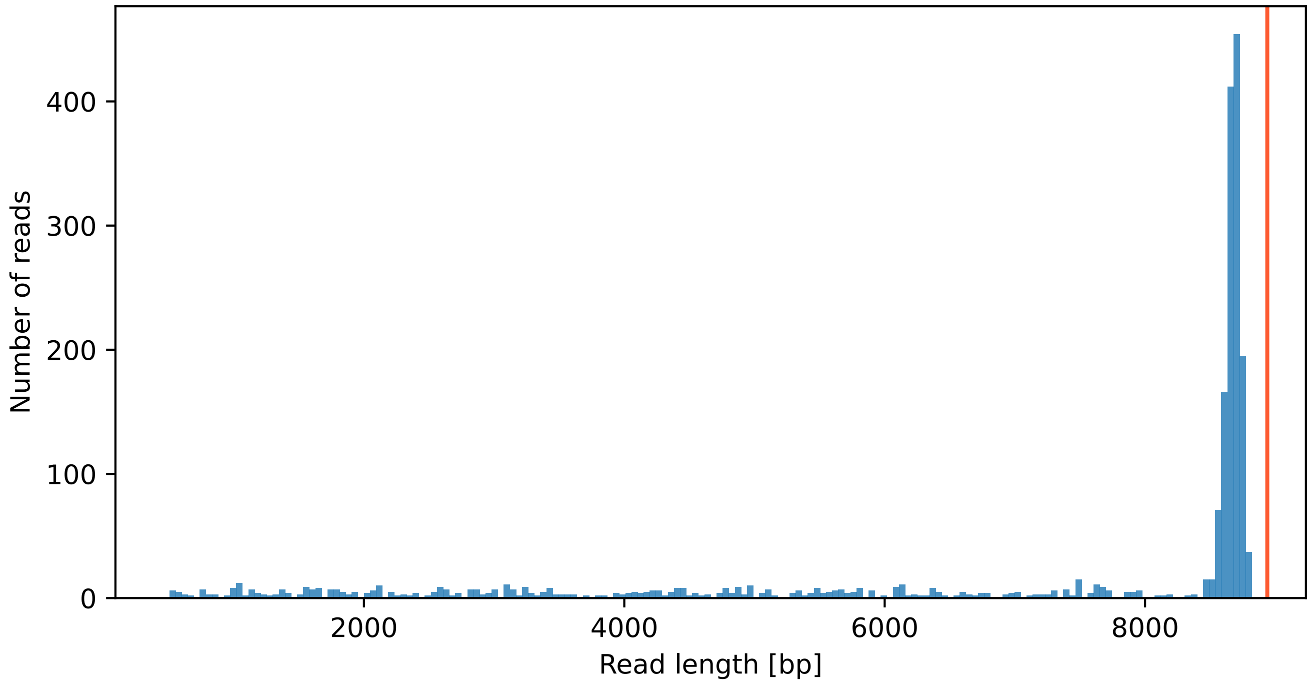


The vertical red lines show the expected construct lengths.

### ☒ EGF2_13

Result: FAIL

Length of reference: **8939** bp.

#### Coverage plot

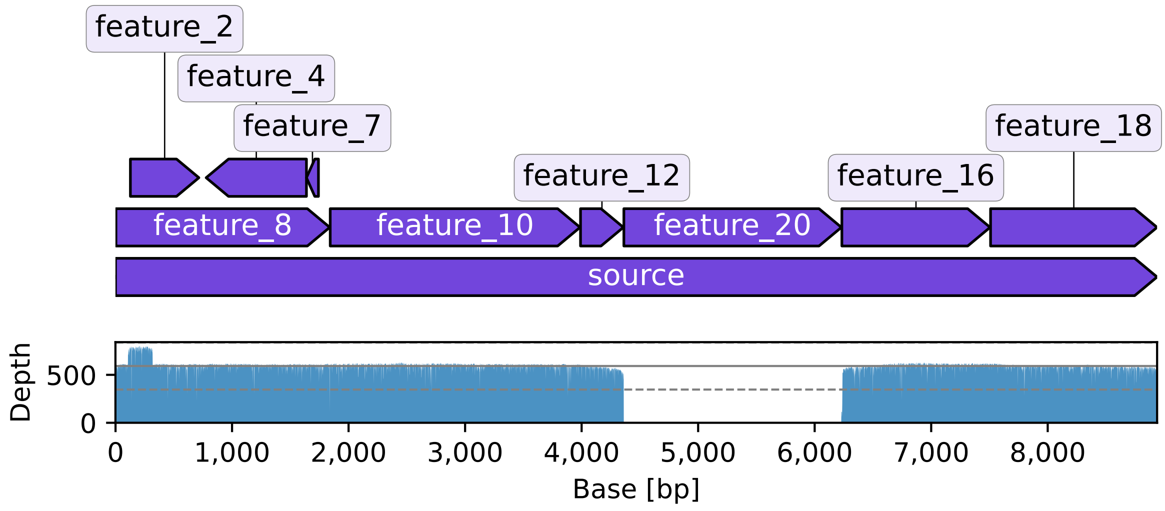


Positions with zero coverage: 4376-6227

Positions with low coverage (<50%): 137, 448, 695, 1837, 3882, 4362-6237, 6499, 6748, 7800

Cumulative plot of longest unaligned interval in each read (81% above cutoff. mode: 1656 bp):

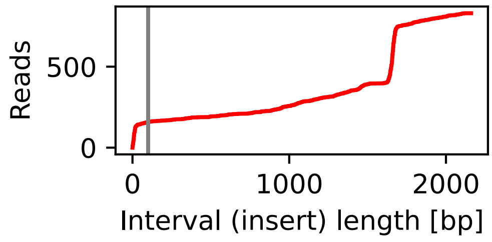


#### Comparison with reference

| LOC | REF | ALT | TYPE | DP | RO | AO | T |
|---|---|---|---|---|---|---|---|
| **1836** | TCA | TA | del | 140 | 1 | 136 | 1 |
| **6233** | TCCCCCCCCCCTAAC... | TCCCCCCCCTAACG | del | 31 | 3 | 15 | 0 |

**EGF2_13** reference vs consensus of reads:

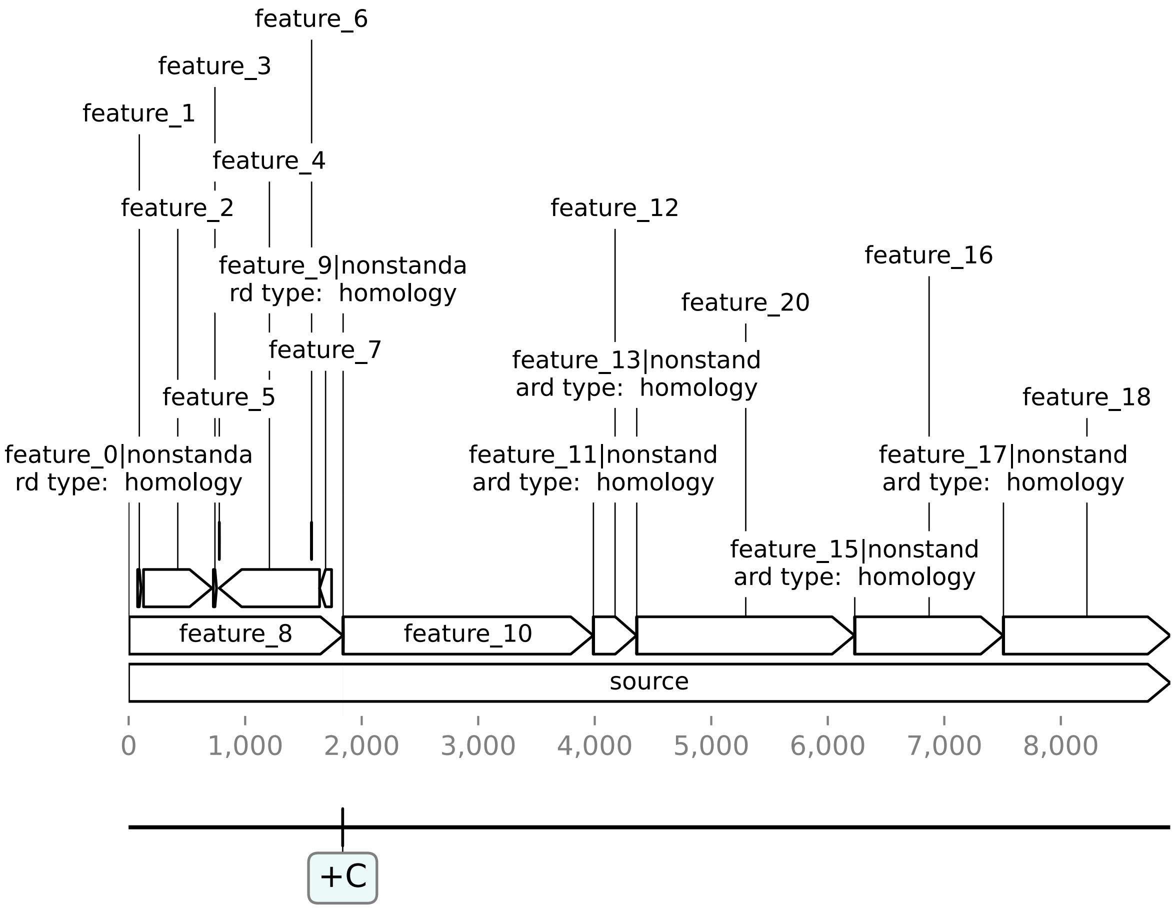

### Appendix

Each barcode chapter consists of a summary and analysis sections.

#### Summary page

The first page summarises the results and other details for each plasmid construct.

#### Analysis pages

Each plasmid construct is analysed separately. The result is summarised with a symbol:
☑ / : correct construct, based on no detection of errors
⚠ / : warning signs are present, review the results and make a decision
☒ / : errors were detected in the construct
⍰ / : sequencing problems, insufficient reads

Depending on settings and input data, some of the described plots may not be in the report.

**Coverage**

The coverage plot displays the number of reads aligning to each base of the reference, in blue. This is useful for detecting deletions. The grey line shows the median. The dashed lines show the ±stdev. Positions with low coverage are also listed, if there are any. Low coverage = below the coverage threshold.

**Comparison with reference**

If a consensus (or *de novo* assembly) sequence is created from the reads, then it is compared with the reference. If the lengths differ more than 5%, then an error is displayed here, otherwise a GeneBlocks plot is made. In this plot, the reference is displayed, and annotated with changes compared to the consensus obtained from the reads (unless a note tells otherwise). For example, +G means that the reference file has an extra G compared to the DNA sample, in other words, the DNA has a G deletion. A VCF (variant call format) table is also displayed. Both the reported and the VCF table positions are zero-based, but the VCF table reports the position of the variation, rather than the nucleotide. Note that the reported depths (DP) may be lower than the ones in the coverage plot, depending on how the variant call was performed. The columns of the table are:

- LOC: 0-based position index (where the first nucleotide has index 0)
- REF: Reference sequence
- ALT: Alternative sequence
- TYPE: The type of allele (either snp, mnp, ins, del or complex)
- DP: Total read depth at the locus
- RO: Reference allele observation count
- AO: Alternate allele observations

An additional column T is provided to mark entries (1) that are deemed true mutations. Inconclusive mutations at repeats (homopolymers) were shown to be systematic sequencing errors, and can be ignored.

**Ediacara**

The report was generated by Ediacara, a software published by the Edinburgh Genome Foundry (EGF). Ediacara is part of the EGF Codons engineering biology software suite for DNA design, manufacturing and validation.
