## Supporting information S2 for "Biofoundry-scale DNA assembly validation using cost-effective high-throughput long read sequencing"

Edinburgh Genome Foundry

### Assembly analysis report

This document reports the sequencing results of the **EGF demo review** project. Each chapter details the results of a plasmid (assembly). Please see the Appendix on the last page for an explanation of the report.

### EGF2_2

**Note**: the assembly is in reverse complement, compared to the reference.

Plot of the aligning parts:

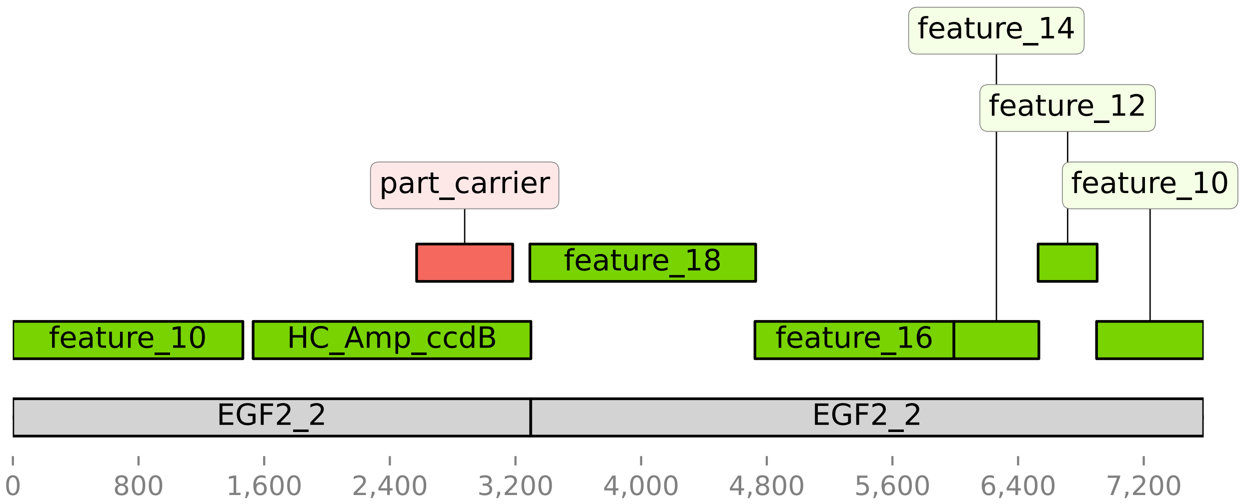


Alignment table:

| Name | Length | Start | End | Strand | T Start | T End | Matches | Size | Quality |
|---|---|---|---|---|---|---|---|---|---|
| **EGF2_2** | 7604 | 3307 | 7604 | - | 3297 | 7582 | 4285 | 4297 | 60 |
| **EGF2_2** | 7604 | 0 | 3307 | - | 0 | 3297 | 3297 | 3307 | 60 |
| **HC_Amp_ccdB** | 2721 | 947 | 2721 | - | 1530 | 3299 | 1769 | 1774 | 60 |
| **feature_10** | 2166 | 6 | 1475 | - | 0 | 1465 | 1465 | 1469 | 60 |
| **feature_10** | 2166 | 1475 | 2159 | - | 6900 | 7582 | 682 | 684 | 60 |
| **feature_12** | 390 | 7 | 383 | - | 6528 | 6904 | 376 | 376 | 60 |
| **feature_14** | 554 | 6 | 547 | - | 5992 | 6533 | 541 | 541 | 60 |
| **feature_16** | 1294 | 15 | 1287 | - | 4725 | 5992 | 1267 | 1272 | 60 |
| **feature_18** | 1451 | 6 | 1445 | - | 3292 | 4730 | 1438 | 1439 | 60 |
| **part_carrier** | 2329 | 235 | 852 | + | 2571 | 3183 | 611 | 617 | 60 |

### EGF2_13

**Note**: the assembly is in reverse complement, compared to the reference.

Plot of the aligning parts:

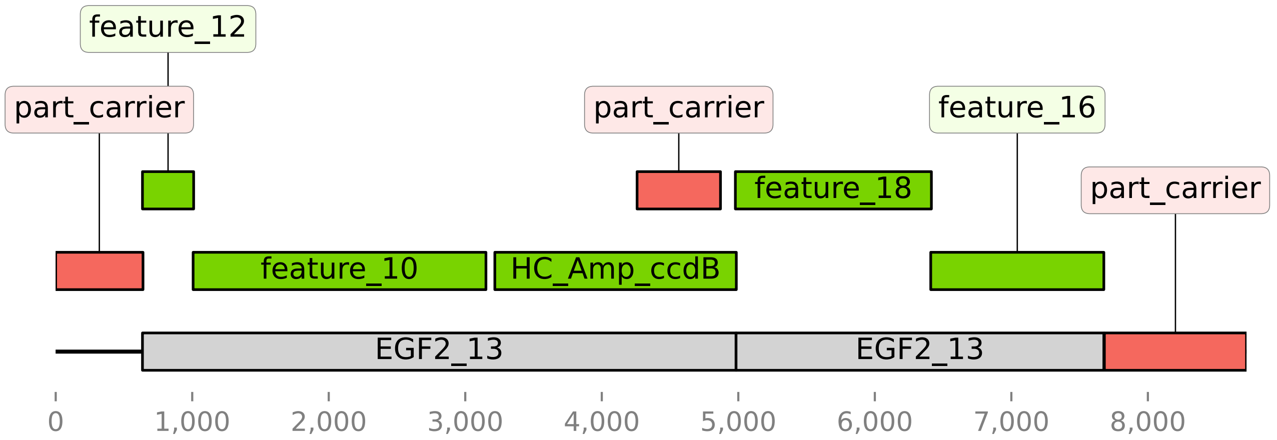


Alignment table:

| Name | Length | Start | End | Strand | T Start | T End | Matches | Size | Quality |
|---|---|---|---|---|---|---|---|---|---|
| **EGF2_13** | 8939 | 0 | 4362 | - | 636 | 4984 | 4348 | 4362 | 60 |
| **EGF2_13** | 8939 | 6238 | 8939 | - | 4984 | 7678 | 2694 | 2701 | 60 |
| **HC_Amp_ccdB** | 2721 | 947 | 2721 | - | 3217 | 4986 | 1769 | 1774 | 60 |
| **feature_10** | 2166 | 6 | 2159 | - | 1007 | 3152 | 2145 | 2153 | 60 |
| **feature_12** | 390 | 7 | 382 | - | 636 | 1011 | 375 | 375 | 60 |
| **feature_16** | 1294 | 15 | 1287 | - | 6410 | 7678 | 1268 | 1272 | 60 |
| **feature_18** | 1451 | 6 | 1445 | - | 4979 | 6415 | 1436 | 1439 | 60 |
| **part_carrier** | 2329 | 1287 | 2329 | - | 7682 | 8723 | 1041 | 1042 | 60 |
| **part_carrier** | 2329 | 646 | 1287 | - | 0 | 640 | 640 | 641 | 60 |
| **part_carrier** | 2329 | 235 | 852 | + | 4259 | 4870 | 610 | 617 | 60 |

### Appendix

Each chapter describes results on a plasmid sequence (*de novo* assembly). The provided (part) sequences were aligned against the sequence assembled from the reads. The plot shows alignment regions as annotations. If there are unannotated segments, then none of the parts aligned there. If an assembly plan is provided, then annotations are coloured based on whether they are expected in the plasmid: green: part is expected in the construct; red: part shouldn't be in the construct. Grey is the reference expected sequence for the plasmid. The alignments are provided in a PAF (pairwise mapping format) table. The columns of the table are:

- Name: Query sequence name
- Length: Query sequence length
- Start: Query start coordinate
- End: Query end coordinate
- Strand: \`+\` if query and target on the same strand; \`–\` if opposite
- T Start: Target start coordinate on the original strand
- T End: Target end coordinate on the original strand
- Matches: Number of matching bases in the mapping (includes mismatches but not indels)
- Size: Number bases, including gaps (indels), in the mapping
- Quality: Mapping quality (0–255 with 255 for missing)
